# RelB represses miR-193a-5p expression to promote the phenotypic transformation of vascular smooth muscle cells in aortic aneurysm

**DOI:** 10.1101/2023.02.21.529372

**Authors:** Yisi Liu, Xiaoxiang Tian, Dan Liu, Xiaolin Zhang, Chenghui Yan, Yaling Han

## Abstract

Aortic aneurysm (AA) is a potentially fatal disease with the possibility of rupture, causing high mortality rates with no effective drugs for the treatment of AA. The mechanism of AA, as well as its therapeutic potential to inhibit aneurysm expansion, has been minimally explored. Small non-coding RNA (miRNAs and miRs) is emerging as a new fundamental regulator of gene expression. This study aimed to explore the role and mechanism of miR-193a-5p in abdominal aortic aneurysms (AAA). In AAA vascular tissue and Angiotensin II (Ang II)-treated vascular smooth muscle cells (VSMCs), the expression of miR-193a-5 was determined using real-time quantitative PCR (RT-qPCR). Western blotting was used to detect the effects of miR-193a-5p on PCNA, CCND1, CCNE1, and CXCR4. To detect the effect of miR-193a-5p on the proliferation and migration of VSMCs, CCK-8, and EdU immunostaining, flow cytometry, wound healing, and Transwell Chamber analysis were performed. In vitro results suggest that overexpression of miR-193a-5p inhibited the proliferation and migration of VSMCs, and its inhibition aggravated their proliferation and migration. In VSMCs, miR-193a-5p mediated proliferation by regulating CCNE1 and CCND1 genes and migration by regulating CXCR4. Further, in the Ang II-induced abdominal aorta of mice, the expression of miR-193a-5p was reduced and significantly downregulated in the serum of patients with aortic aneurysm (AA). In vitro studies confirmed that Ang II-induced downregulation of miR-193a-5p in VSMCs by upregulation of the expression of the transcriptional repressor RelB in the promoter region. This study may provide new intervention targets for the prevention and treatment of AA.

## 1. Introduction

Aortic aneurysm (AA) is characterized by the degradation of the aortic wall matrix with the activation of matrix metalloproteinases (MMPs) and inflammation. Inflammation is one of the leading causes of cardiovascular death, as clinical treatments are limited [1–3]. Vascular smooth muscle cells (VSMCs) are the main components of the vascular wall and play an important role in maintaining the vascular structure. The phenotypic transformation of VSMCs is a critical pathological factor in AA that manifests as dedifferentiation, proliferation, migration, and an increase in matrix metalloproteinases. In contrast, inhibiting VSMC phenotypic transformation attenuates AA progression [4]. Therefore, understanding the mechanisms and new targets involved in the phenotypic transformation of VSMCs and developing new drugs are the key strategies to prevent and treat AA.

MicroRNAs are small non-coding RNAs that bind to the 3’-UTR of the target gene and inhibit translation [5]. Evidence suggests that miRNAs participate in the occurrence and development of AA by regulating the phenotypic switch of VSMCs. Our preliminary results obtained from an Angiotensin II (Ang II)-induced mouse model using microRNA chip analysis suggest the presence of 65 differentially expressed miRNAs in normal blood vessels and AA (Figure S1A-S1C). In the AA group, the expression of miR-193a-5p was downregulated by 6.25 times compared to that in the control group. Several studies have reported a significant reduction in the expression of miR-193a-5p in the serum or vascular tissue of patients with AA or aortic dissection [6–8]. MiR-193a-5p is a member of the miR-193 family and is located on human chromosome 17q11.2. A recent report showed that miR-193a-5p can downregulate the expression of target genes by binding to the 3’-untranslated region (UTR) of the target gene mRNA, such as CCNE1 or CCND1, and regulate tumor cell proliferation, migration, and invasion [9–11]. However, the mechanism of action is yet to be elucidated. In the present study, we observed the regulatory effect of miR-193a-5p on the phenotypic transformation of muscle cells and explored the potential molecular mechanism involved in its downregulation in AA vascular tissue.

## 2. Material and methods

### Patients

Serum samples were collected from 8 healthy subjects (40±10years old, 6 cases male) and 4 patients with abdominal aortic aneurysm (50±10years old, 3 cases male) in Northern Theater Command General Hospital. Serum was stored at −80 °C Patients enrolled in this study did not have valvular heart disease, chronic kidney disease, autoimmune disease or other cardiopulmonary organic diseases. This research was in line with the World Medical Association Code of Ethics (Declaration of Helsinki) (Lun Review No. Y (2012)002). This study was approved by the Ethics Committee of Northern Theater General Hospital. All enrolled patients signed informed consent prior to the study.

### Cell Culture

Primary mouse vascular smooth muscle cells were isolated and cultured in medium containing 10% fetal bovine serum (FBS) and 2% antibiotic DMEM. In briefly, VSMCs were isolated from the aortas of 10-week-old C57BL/6 mice. The aortas were excised, washed in phosphate buffered saline, and incubated in DMEM containing 1 mg/mL of Collagenase type II (Sigma) for 10 to 15 minutes. Then, the adventitia was removed, and the endothelial cells were gently scraped off. The vessels were transferred to culture dishes containing DMEM with 1 mg/mL of Collagenase type I (Sigma) and 0.125 mg/mL of Elastase type III (Sigma), minced with scissors, incubated in 37°C, 5% CO2 humidified incubator until 90% of the cells were dispersed under the microscope. The cells were centrifuged at 1600 rpm for 5 minutes, then resuspended in 3 mL DMEM with 10% fetal calf serum (FCS), 2% penicillin-streptomycin, and cultured in plates or flasks.

### Cell treatment

When the cell density reached 80%, the cells were treated with Ang II and collected 24 hours later.

### Extraction and culture of primary vascular smooth muscle cells from mice

The aorta of mice was separated, cut into pieces with scissors, and beaten with DMEM. After centrifugation for 5min, DMEM was discarded, and the aortic tissue was planted in the petri dish, collagenase was added for digestion for 2h. If adhered to the wall, the culture medium was changed to the DMEM containing 20% fetal bovine serum (FBS), and the solution was changed every day. Intervention could be given when the cells reached 6-8 generations.

### Cell Transfection

miR-193a-5p mimic and mimic NC, inhibitor, inhibitor NC and siRNA for CCND1, CCNE1 and CXCR4 were purchased from Ribobio. pcDNA3.1-CCND1, CCNE1 and CXCR4 vectors were purchased from GENEWIZ. Mimic, inhibitor, siRNAs and plasmids were transfected into cells with RNA IMAX or Lipofectamine 2000 (Invitrogen).

### Western Blot

Cells were collected by RIPA (Thermo scientific) ice lysis for 30 min, centrifuge at 4°C (Heraeus, Germany) 12,000 rpm for 15 min, supernatant was taken, protein concentration was measured by BCA (Thermo scientific). Sample 20 μg denatured protein 10% sodium dodecyl sulfate polyacrylamide gel electrophoresis. After the protein gel was transferred to the polyvinylidene fluoride (PVDF) membrane, incubation in a closed solution containing 5% fatless milk powder, then 1:1000 diluted primary antibody (PCNA, CCND1, CCNE1, α- SMA and CXCR4) was incubated at 4°C overnight. After the membrane was washed with TBST, the secondary antibody was diluted at 1:1000 and incubated at room temperature for 1h. ECL (GE) luminescence developed to detect the expression of protein. For the extraction of vascular tissue protein, the vascular tissue was added to RIPA and crushed in the grinder (Servicebio) and the protein could be obtained by centrifuge at 4 °C. The remaining steps are the same as for cell extraction.

### Quantitative Real-time PCR

RNA from cells or serum was extracted by Trizol reagent (Thermo), and cDNA was retro-transcribed by miDETECT A Track miRNA qRT-PCR Starter Kit (Ribobio). qRT-PCR was used to detect the expression of miR-193a-5p. U6 was used as an internal reference. Similarly, miR-39 was used as an internal reference for serum. mRNA was retro-transcribed by Takara kit (Takara Bio), GAPDH was used as internal reference. For RNA extraction from vascular tissue, Trizol is added to vascular tissue and crushed in the grinder (Servicebio). The remaining steps are the same as for cell extraction.

### Cell Counting Kit-8 (CCK-8) Assay

CCK8 kit (Beyotime) was used to detect the proliferation activity of VSMCs. VSMCs were seeded into 96-well plates, and 10 μL CCK8 solution was added to each well 24h after adherence. Absorbance was detected at 450nm with a microplate for 1h.

### EdU cell proliferation assay

The proliferation of VSMCs was detected by EdU kit (Ribobio). In brief, the VSMCs were planted into the 24-well plate changed into 50 μM EdU medium 300 μL (10%DMEM 1:1000 dilution) and incubated for 2 h. The cells were washed with PBS twice, then 500 μL paraformaldehyde fixative (Servicebio) was added to each well to fix the cells for 30 min. 300 μL 2 mg/mL glycine for 5 min. Add 300 μL PBS wash for 5 min, add 300 μL 0.5% Triton x-100 (Sangon Biotech) to penetrant cells for 10 min, wash with PBS once. 300 μL 1xApollo staining solution for 30 min at room temperature. Hoest4432 solution was diluted with 100:1 deionized water, and 300 μL was added to each well for 30 min without light. The tablets were photographed under an inverted microscope.

### Cell cycle detection by flow cytometry (PI method)

Flow cytometry was used to measure the cell cycle of VSMCs and sheath fluid was prepared one day in advance. The cells were digested with trypsin and centrifuged at 1000 rpm for 10 min. Wash PBS twice. The cells were re-suspended with 0.6 mL PBS and transferred to glass tubes. Added 1μl PI dye, filtered with 300 mesh, and checked on the machine.

### Invasion Assay

Transwell chamber (24-well plate, 8.0 μm COSt Qr LOT 25820052) was used to detect VSMC invasion. 200 μL serum-free cell suspension was added in the upper chamber. The lower compartment was incubated with 500 μL 10% serum FBS DMEM (Gibco). 24 h later, the lower compartment was washed twice with PBS (Sangon Biotech) and fixed with 1mL paraformaldehyde solution (Servicebio) for 30 min. Then the lower compartment was stained with Giemsa stain A: B (1:1) for 30 min and A: B (1:3) for 10 min. Photograph with light microscope (Carl Zeiss AG)

### Wound healing Assay

The VSMCs were incubated into the six-well plate with 80% fusion. A cross was drawn longitudinally and laterally with 1μL small spear tip. The initial conditions at 0 h and migration at 24 h were recorded with an inverted microscope (Olympus Corporation).

### Prediction of miRNA target genes or transcriptional factor

Using bioinformatics methods to predict the target genes that differential miRNAs may bind to, we also analyzed the target genes that may be regulated by up regulating differential miRNAs and down regulating differential miRNAs respectively. The databases involved are as follows: miRWalk, miRDB. Finally, obtain the intersection genes of the two databases for subsequent analysis.

To prediction of transcriptional factor of miRNA, we searched for TFs in the Jasper Database that may regulate miR-193a-5p and screened for its preferred homology of binding site and superior ranking grade in the Jasper Database.

### Luciferase reporter gene

The construction of CXCR4 3’UTR vector, mir-193a-5p promoter, PCDNA3.1Relb and pcDNA3.1 C/EBPβ were purchased from GENEWIZ. CXCR4 3’UTR was co-transfected with mir-193a-5p mimic by HEK293 cells, and mir-193a-5p promotor was co-transfected with pcDNA3.1Relb and pcDNA3.1 C/EBPβ for 48h. Luciferase activity was detected.

### Immunofluorescence

Slides were laid, 24h adherent was washed with PBS 3 times, 500μL paraformaldehyde (Servicebio) fixative solution was fixed for 15min, 0.2% TritonX-100 (Sangon) permeable cells were prepared for 10min, washed with PBS twice, and goat serum was soaked and sealed at room temperature for 30min. After that, PBS was diluted with 1:100 primary antibody (CCND1, CXCR4) at 4 °C overnight, and rewarmed for 30min the next day. Fluorescence secondary antibody was prepared (diluted with PBS at 1:200), and incubated for 1h in darkness, washed with PBS for 30min, and nucleated with DAPI for 5min, washed with PBS for 3 times. The tablets were sealed with anti-fluorescence quenching agent and photographed by optical microscope (Carl Zeiss AG)

### Statistical Analysis

The above experiments were repeated three times, and Prism8 software was used for statistical analysis. T-test and one-way ANOVA were used. P <0.05 was considered statistically significant.

## 3. Results

### 3.1. MiR-193a-5p inhibits the proliferation and migration of VSMCs in vitro

As shown in Figures S2A and S2B, miR-193a-5p expression increased in mouse vascular tissue and vascular smooth muscle cells but was reduced in the serum of AA patients compared to that in control patients (Figure S2C). Furthermore, miR-193a-5p expression decreased significantly in Ang II-stimulated primary mouse VSMCs in a dose-dependent manner (Figures S2D and S2E).

To determine the effect of miR-193a-5p on Ang II-stimulated VSMCs impairment, primary mouse VSMCs were transfected with mimic NC or mimic miR-193a-5p. The miR-193a-5p expression increased significantly in the cells transfected with mimic miR-193a-5p compared to cells transfected with mimic NC (Figure 1A). CCK8, EdU staining, and immunoblotting for cell cycle proteins PCNA and cyclin D1 revealed that overexpression of miR-193a-5p suppressed Ang II (1μM)-induced proliferation of VSMCs (Fig. 1B-1F). Transwell chamber assays and wound-healing tests revealed that an increase in miR-193a-5p remarkably blocked Ang II-induced migration of VSMCs when compared to mimic NC-transfected cells (Figure 1G–1J). These results suggest that the overexpression of miR-193a-5p inhibited Ang II-induced proliferation and migration of VSMCs. However, transfection with a miR-193a-5p inhibitor or NC inhibitor for 24h increased the Ang II-induced proliferation and migration of VSMCs (Figures 2A–2H). Interestingly, miR-193a-5p had no regulatory effect on VSMC differentiation as measured by SM α-actin expression in cells treated with miR-193a-5p mimic or control (Figure S3A and S3B).

**Figure 1.**
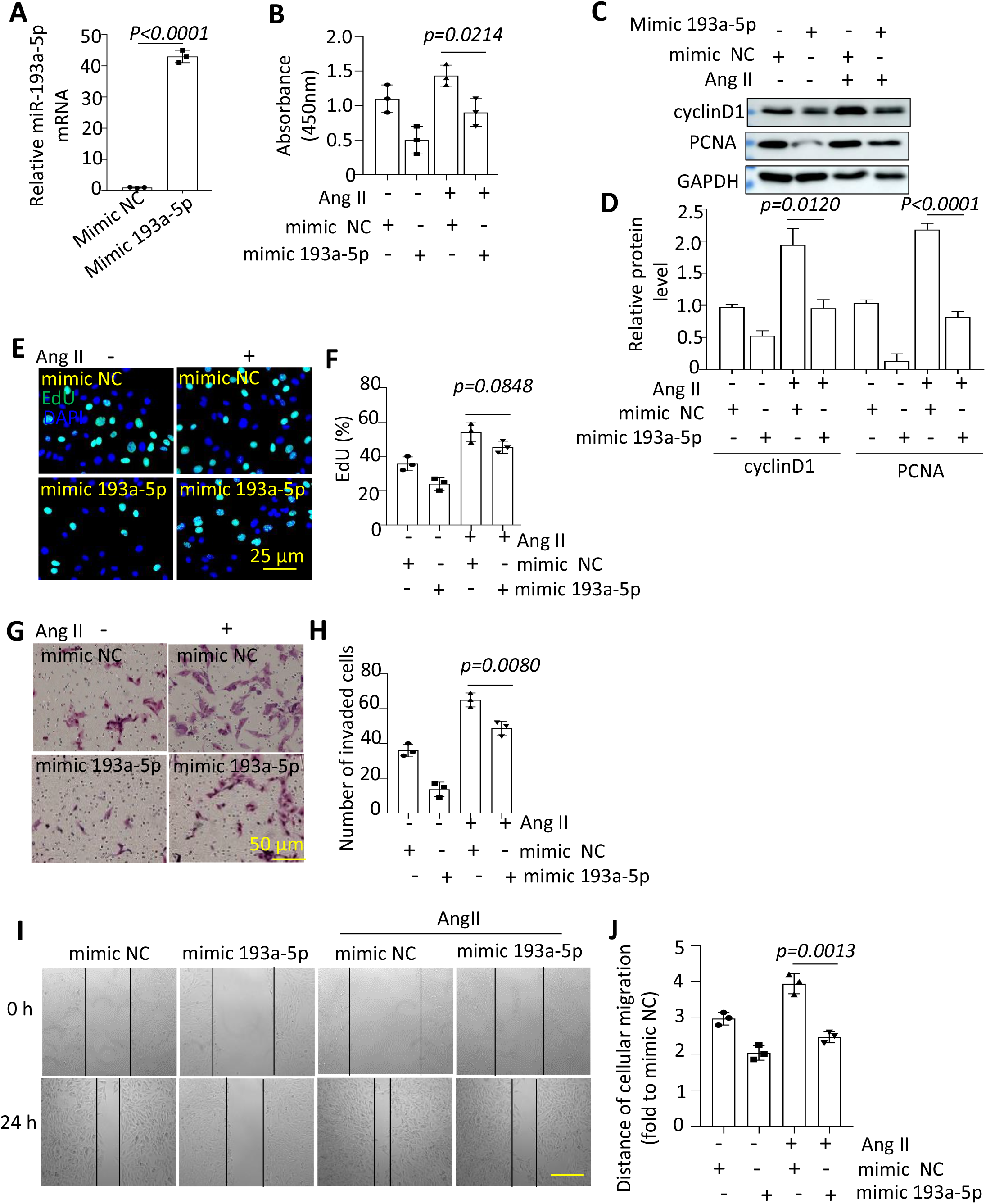
Mir-193a-5p could inhibit the proliferation, migration, and invasion of vascular smooth muscle cells (VSMCs). (A) VSMCs were transfected with mimic or mimic control expressing miR-193a-5p. The transfection efficiency of mir-193a-5p was detected by qRT-PCR. (B) Proliferation activity was detected by CCK8 assay. To induce the proliferation and migration, cells were stimulated with Ang II (1 μM) for 24 h. (C-D) Western Blot was used to detect the protein levels of PCNA and cyclinD1. (E-F) EdU staining imaging was used to detect proliferation. (G-H) Transwell invasion assay was used to detect migration. (I-J) The migration was observed by wound healing assay.

**Figure 2.**
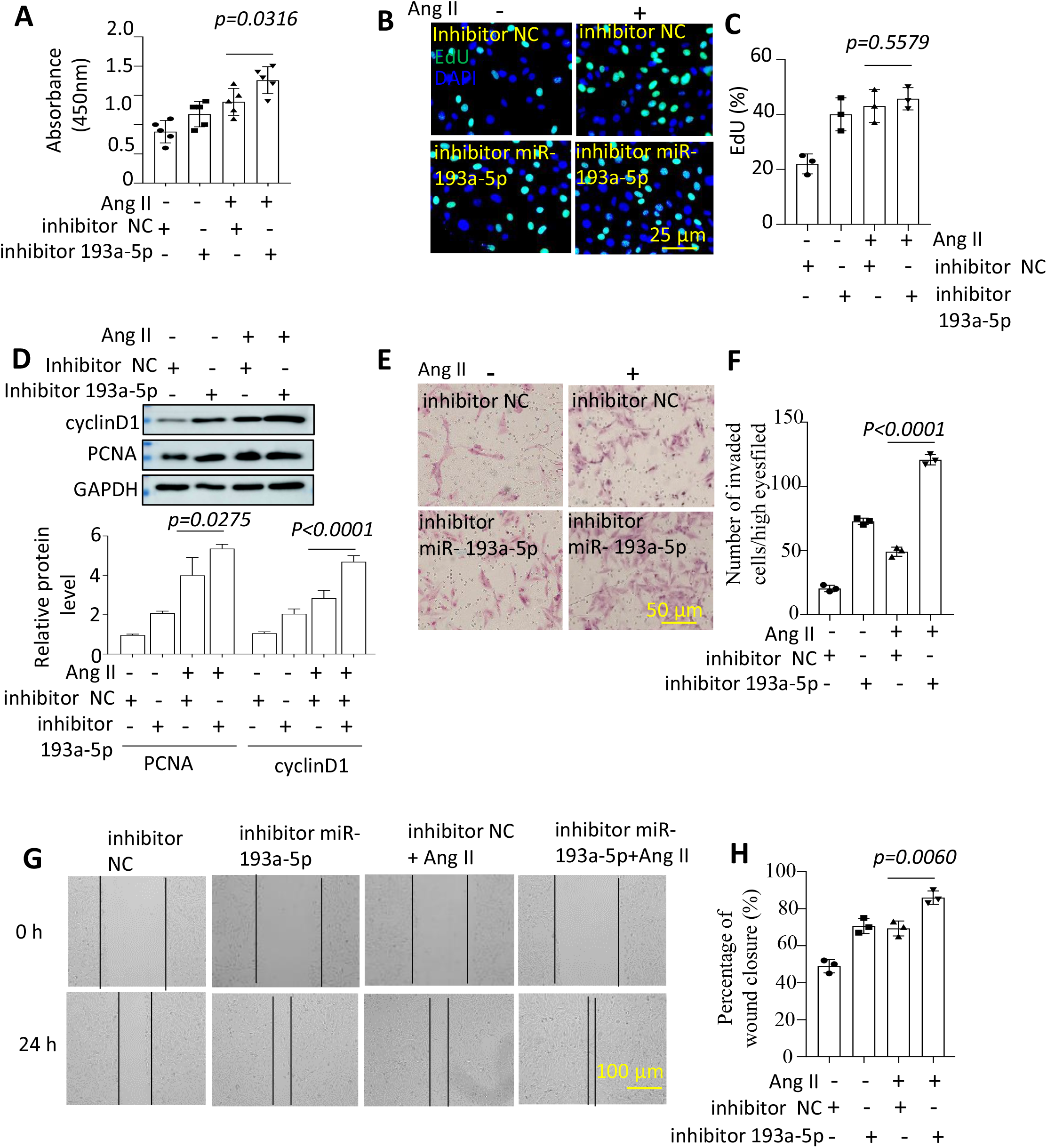
Inhibition of mir-193a-5p promotes the proliferation and migration of VSMCs. Vascular smooth muscle cells were transfected with inhibitor mir-193a-5p for 24h and then stimulated with Ang II (1 μM) for 24 h. (A-C) CCK8 assay and EdU staining imaging were used to detect proliferation. (D) Western blotting was used to detect protein levels of PCNA, and cyclinD1. (E-F) Transwell invasion assay was used to detect cell migration. (G-H) Migration was observed using a wound-healing assay.

### 3.2. MiR-193a-5p modulated the expression of CCND1, CCNE1 and CXCR4

To elucidate the mechanisms involved in the VSMC phenotype switching, we analyzed the downstream target genes regulated by miR-193a-5p. TargetScan, miRDB, and PicTar software predicted that the 3’UTR regions of CCND1, CCNE1, and CXCR4 may interact with miR-193a-5p and act as the binding sites (Figure 3A). Furthermore, an increase in miR-193a-5p inhibited the expression of CCND1, CCNE1, and CXCR4 at both the mRNA and protein levels in VSMCs, and the downregulation of miR-193a-5p increased the expression of CCND1, CCNE1, and CXCR4 significantly (Figure 3B to 3G). Similarly, immunofluorescence staining and quantification revealed that the miR-193a-5p mimic decreased CCND1 and CXCR4 expression, while the inhibitor of miR-193-5p increased both CCND1 and CXCR4 expression in VSMCs (Figures 3H and 3I). Since CCNE1 and CCND1 are downstream regulators of miR-193a-5p, we further examined the interaction of CXCR4 with miR-193a-5p. Luciferase fluorescent reporter gene analysis confirmed that miR-193a-5p regulates the expression of CXCR4 by interacting with its 3’UTR region (Figure 3J).

**Figure 3.**
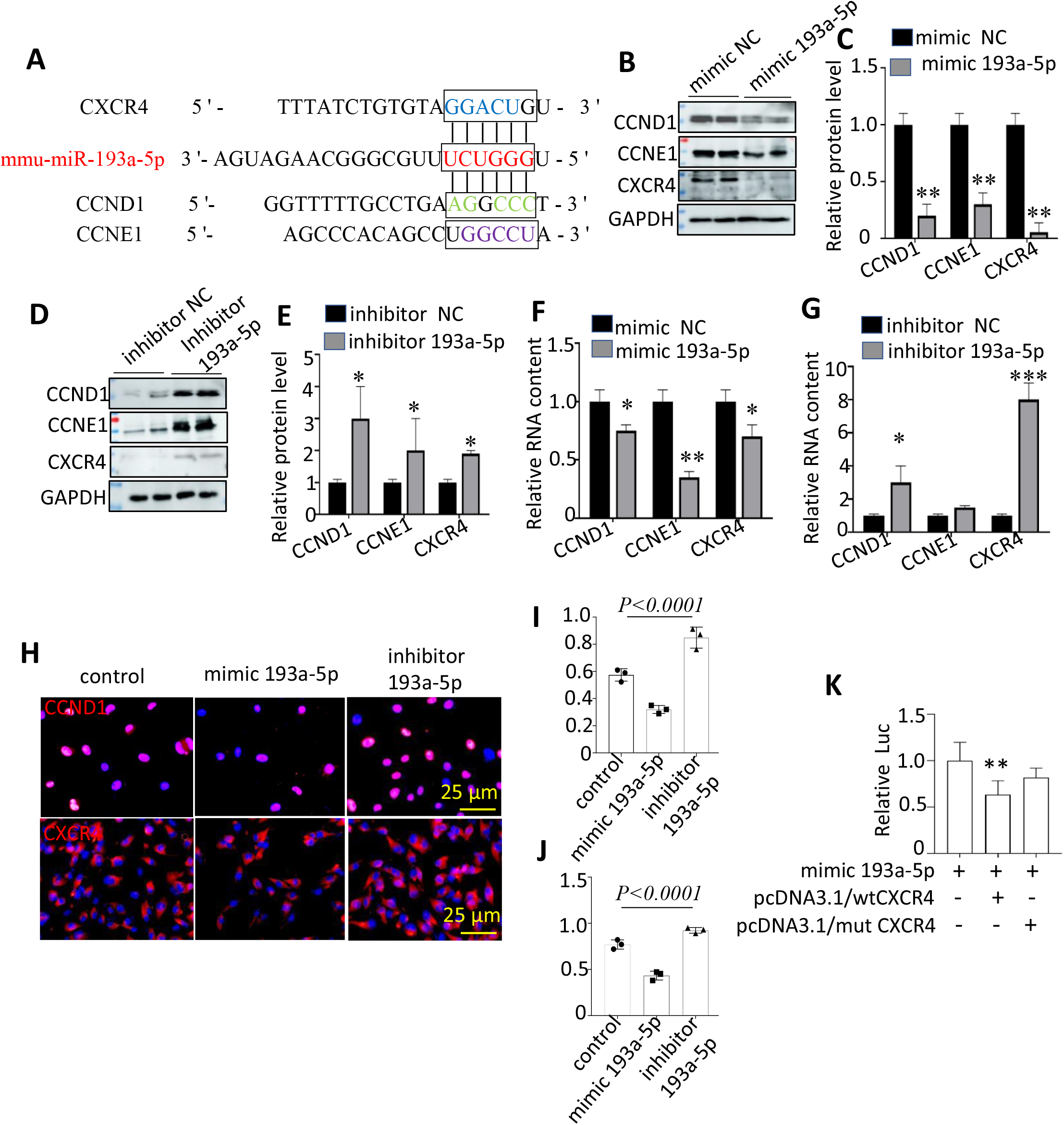
MiR-193a-5p modulated the expression of CCND1, CCNE1, and CXCR4. VSMCs were infected with a mimic or mimic control expressing miR-193a-5p, and 24h later, total RNAs or protein was isolated. (A) Binding sites of mir-193a-5p on CCND1, CCNE1, and CXCR4. (B-E) Western blot was used to detect CCND1, CCNE1, and CXCR4 protein levels. (F-G) Real-time quantitative PCR was used to detect the CCND1 and CCNE1 mRNA levels. (H) Immunofluorescence was used to detect CCND1 and CXCR4 expression. (I) The relationship between mir-193a-5p and CXCR4 was detected using a dual-luciferase reporter gene. \**P* < 0.05, \*\**P*<0.01, \*\*\**P*< 0.001 compared to NC. Luc: luciferase activity; CCND1: CyclinD1, CCNE1: CyclinE1.

### 3.3. CCND1 and CCNE1 are involved in the VSMCs proliferation promoted by the downregulation of miR-193a-5p

To investigate if CCND1, CCNE1, and CXCR4 mediated the phenotypic switch of VSMCs and were regulated by miR-193a-5p, we overexpressed CCND1, CCNE1, and CXCR4 in cells. Interestingly, both CCND1 and CCNE1 overexpression significantly reversed the miR-193a-5p mimic-induced proliferation inhibitory effect (Figures 4A–4I). In contrast, when CCND1 and CCNE1 were silent for 24h, the proliferative potential was inhibited in VSMCs transfected with the miR-193a-5p inhibitor (Figures 5A–5C, and Figure S4A-S4C). Western blotting for PCNA also indicated that CCND1 and CCNE1 mediated the miR-193a-5p-inhibited VSMCs proliferation (Figure 5D–5H). However, as shown in Figure S5A-S5D, silencing or overexpressing CXCR4 did not change the expression of PCNA, suggesting that CXCR4 did not regulate the proliferation of VSMCs.

**Figure 4.**
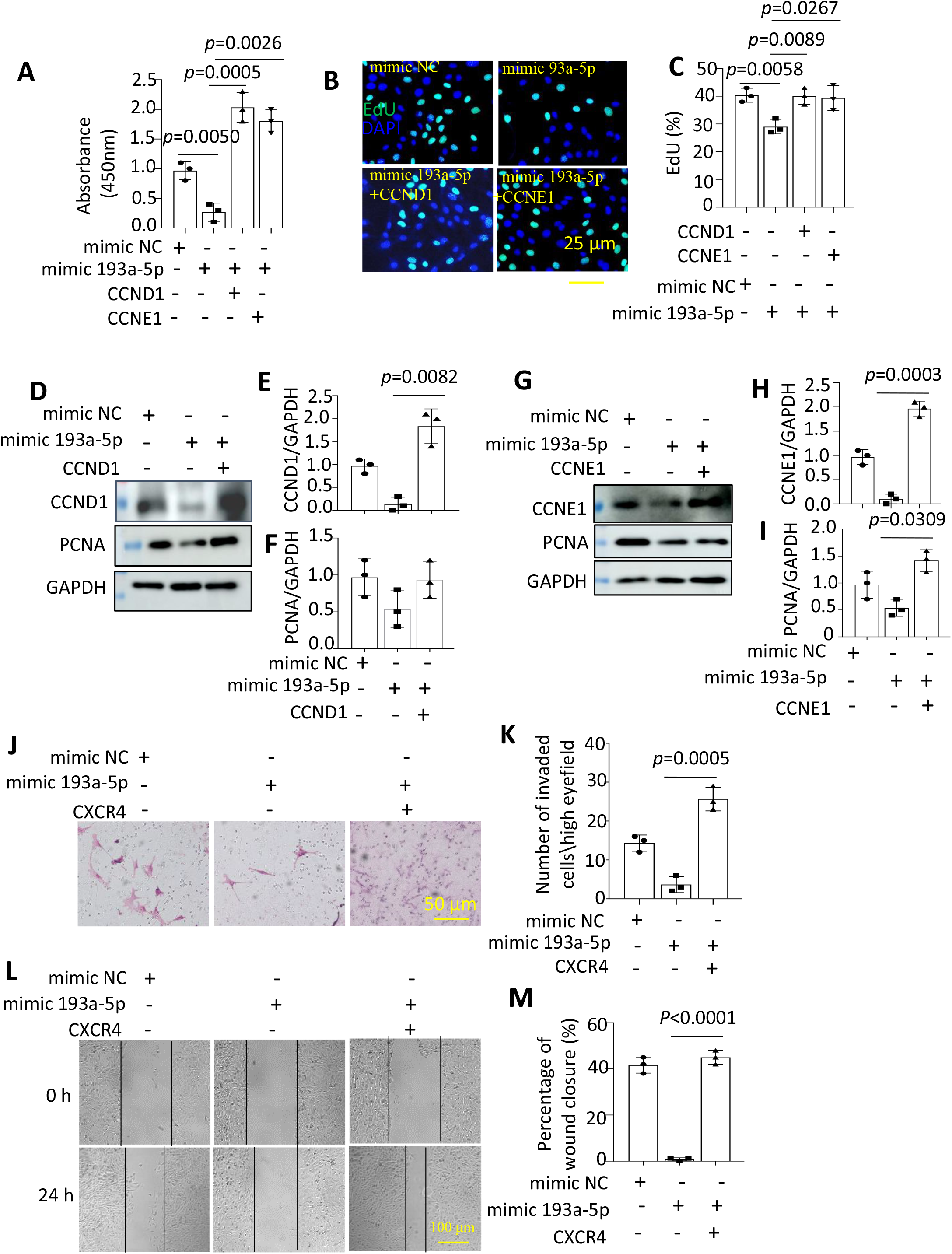
Mir-193a-5p inhibited the proliferation and migration of VSMCs by targeting CCND1, CCNE1, and CXCR4. (A) Proliferation activity was detected using CCK8 assay. (B-C) EdU staining imaging was used to detect proliferation. (D-I) Western Blot was used to detect the protein levels of CCND1, CCNE1, and PCNA. (J-K) Transwell invasion assay was used to detect migration. (L-M) The migration was observed by Wound healing assay. #P>0.05, *P < 0.05, **P<0.01, ***P< 0.001 mimic NC.

**Figure 5.**
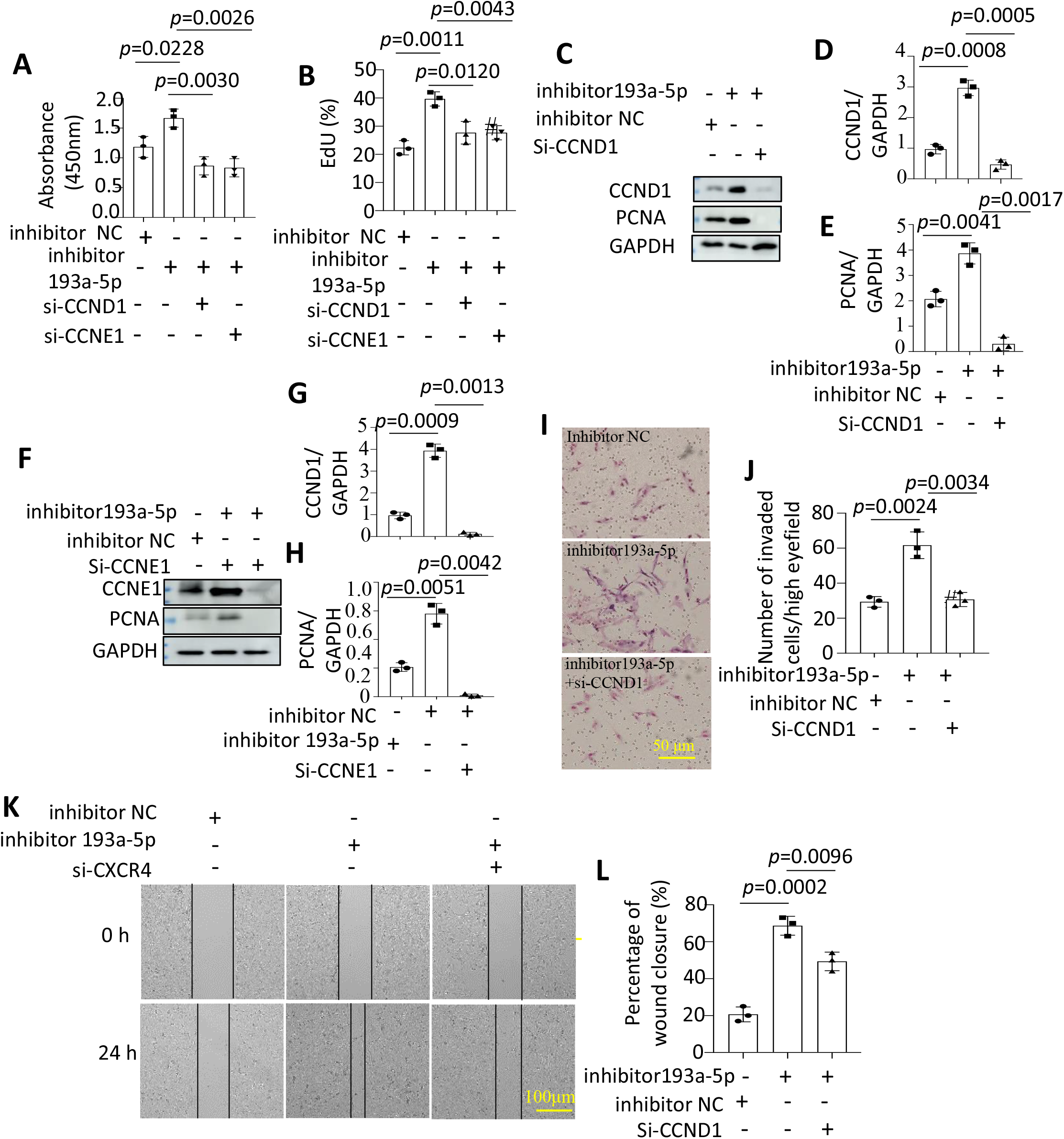
Inhibition of mir-193a-5p increased the proliferation and migration of VSMCs via CCND1, CCNE1, and CXCR4. (A) Proliferation activity was detected using CCK8 assay. (B) EdU staining imaging was used to detect proliferation. (C-H) Western Blot was used to detect the protein levels of CCND1, CCNE1, and PCNA. (I-J) Transwell invasion assay was used to detect migration. (K-L) The migration was observed by Wound healing assay. si-CXCR4(2) was used. #P>0.05, *P < 0.05, **P<0.01, ***P< 0.001 compared to inhibitor NC.

### 3.4. CXCR4 mediated the migration of VSMCs regulated by miR-193a-5p

Similarly, when VSMCs were transfected with pcDNA3.1 CXCR4 plasmid only, the inhibitory effect of miR-193a-5p overexpression significantly reversed VSMC migration (Figures 4J–4M and Figures S6A–S6B). In contrast, silencing CXCR4 for 24 h blocked the migration and invasion induced by the miR-193a-5p inhibitor (Figure 5I–5L). However, silence of CCND1 or CCNE1 for 24h did not change the invasion potential of VSMC. These results implied that CXCR4 mediates miR-193a-5p-induced migration of VSMCs.

### 3.5. RelB regulated the expression of miR-193a-5p in Ang II-treated VSMCs

To elucidate the reduction of miR-193a-5p in Ang II-treated VSMCs, we searched for TFs in the Jasper Database that may regulate miR-193a-5p. Among them, RelB was screened for its preferred homology with the binding site and superior ranking grade in the Jasper Database (Figure 6A). Western blot analysis showed that RelB increased in Ang II-treated VSMCs and that it was associated with an increase in CCND1, CCNE1, and CXCR4 (Figures 6B and 6C). Furthermore, to understand the impact of RelB on VSMCs, we observed if the overexpression of RelB could decrease miR-193a-5p expression or if inhibition of RelB increased miR-193a-5p expression and decreased the expression of CCND1 and CCNE1. CCND1 expression increased slightly when RelB was overexpressed (Figures 6D–6H). Meanwhile, western blotting showed that RelB expression was dramatically increased in AA mice aortic tissue compared to that in control aortic tissue (Figures 6I–6J). Subsequently, we constructed plasmids in the promoter region of miR-193a-5p and expressed luciferase reporter genes using pcDNA3.1-RelB.

**Figure 6.**
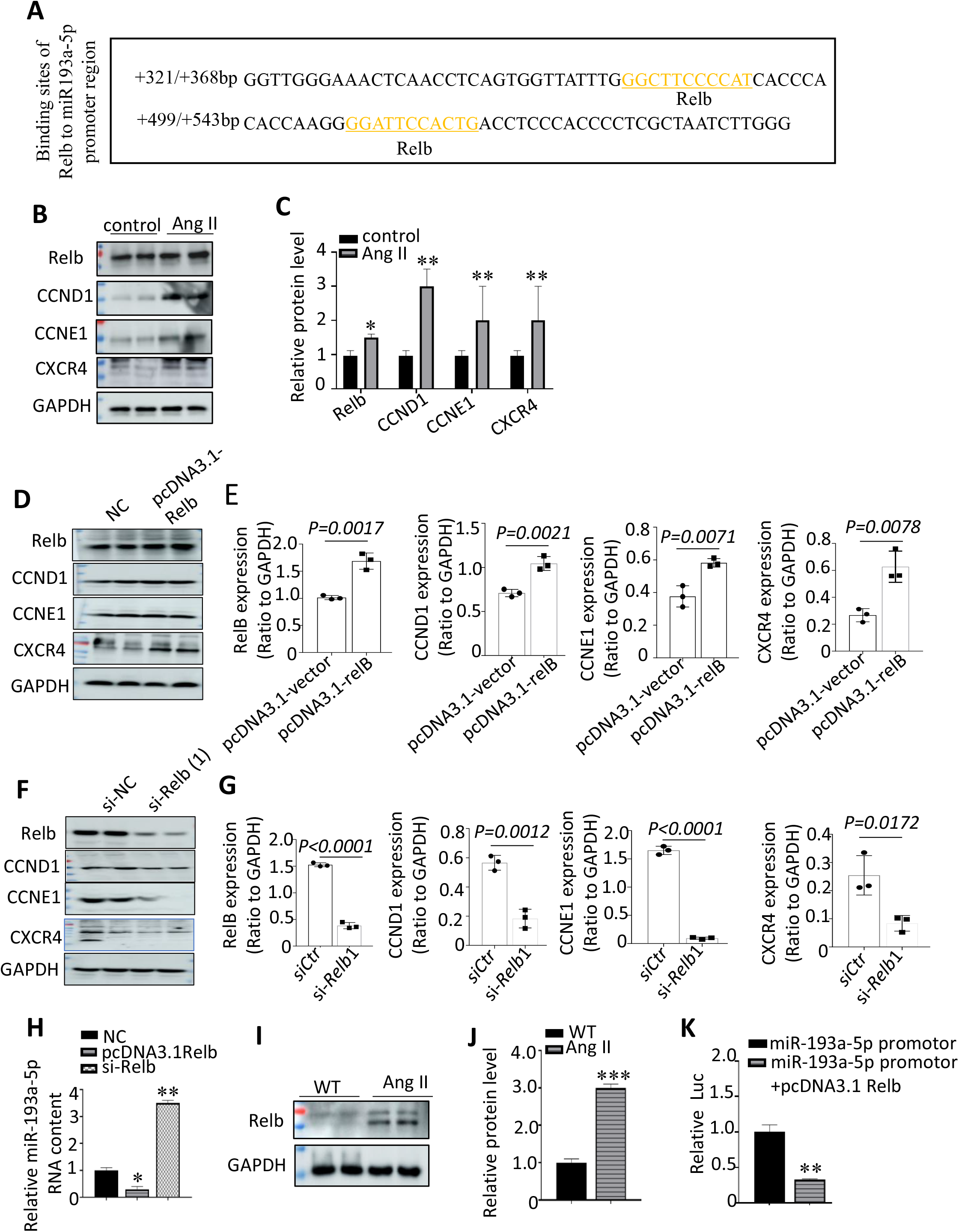
Transcription factor RelB represses the transcription of mir-193a-5p. (A) In the mir-193a-5p promoter region, yellow is the site where RelB may bind to mir-193a-5p. (B-C) Western blot was used to detect the changes in transcription factors and target genes before and after Ang II stimulation. (D-G) Expression changes of mir-193a-5p and CCND1, CCNE1 were detected by interfering and overexpressing RelB. (H) Expression changes of mir-193a-5p were detected by interfering and overexopressing RelB. (I-J) Western blot was used to detect RelB expression in vascular tissues of AAA mice. (K) The relationship between mir-193a-5p and RelB was detected by the luciferase reporter gene. *P < 0.05, **P<0.01, ***P< 0.001 compared to NC. WT: wild-type mouse.

## 4. Discussion

In recent years, there has been an annual increase in the incidence and mortality rates due to AA. However, owing to the lack of clinical manifestations in most AA patients, the patients die once they develop AA. AA can only be detected accidentally in various imaging examinations. Risk factors for AA include age, smoking history, hypertension, and genetic factors. Despite extensive intervention studies targeting these risk factors, including endovascular repair, PPARγ agonists, cell therapy, and kinase inhibitors, no effective clinical treatment has been available until now [12–14].

Many clinical trials and animal models have revealed that the proliferation and migration of VSMCs are critical factors in the pathogenesis of AA [15]. VSMCs have a differentiated phenotype in a resting environment. When stimulated, VSMCs show phenotypic switching and an increase in proliferation and migration, resulting in vascular injury [16]. We propose new therapeutic intervention avenues for the prevention of vascular injury and treatment of AA in order to unravel the regulatory mechanism in VSMC phenotypic transformation.

MicroRNAs are non-coding RNAs that are 17-23 nucleotides long. An increasing number of studies have identified that miRNAs play an important role in the pathogenesis of aneurysms, especially in the regulation of VSMCs [17]. Our preliminary data, consistent with previous studies [6–8] showed that the expression of miR-193a-5p was significantly reduced in mice AA tissue compared to that in control blood vessels. Moreover, miR-193a-5p was highly expressed in primary cultured VSMCs and dramatically downregulated in Ang II-induced VSMCs. This suggests that miR-193a-5p may be involved in the Ang II-induced phenotypic switch of VSMC. In the current study, we confirmed that miR-193a-5p regulates the proliferation and migration of VMSCs by targeting various downstream genes but does not affect cell differentiation. In the present study, we reported that CCND1, CCNE1, and CXCR4 are downstream targets of miR-193a-5p [18,19]. Both CCND1 and CCNE1 are cell cycle regulating genes, and the protein production of cyclin D1 and cyclin E1 is increased in human tumor cells, which regulate the transition of the cell cycle from the G1 restriction point to the S phase, thus playing an important role in the regulation of cell proliferation [20, 21]. CXCR4 is a 352 amino acid rhodopsin-like GPCR and comprises an extracellular N-terminal domain, seven transmembrane helices, and an intracellular C-terminal domain [22]. CXCR4 can exist in the plasma membrane as a monomer, dimer, higher-order oligomer or nanocluster. Cancer cells have increased levels of CXCR4 compared with normal cells [23]. Many studies have revealed that the binding of the CXCR4 receptor to its ligands triggers multiple signaling pathways that orchestrate cell migration, hematopoiesis, cell homing, and retention in the bone marrow. A recent report showed that CXCR4 directly controls the proliferation of non-hematopoietic cells in the presence of the HMGB1·CXCL12 complex. However, in our study, we did not detect the CXCR4 controlling-VSMC proliferation. This phenomenon suggests that CXCR4 regulating cell proliferation is closely related to the cell type[23]. Our study also suggests that CCND1 and CCNE1 are involved in the proliferation of VSMC regulated by miR-193a-5p, while CXCR4 mediates its migration.

RelB, a member of the NF-KB family, is an activated subunit and inflammatory factor. It has been reported that RelB participates in the regulation of VSMCs proliferation, which may be due to the regulation of c-myb gene transcription [24]. Accumulating evidence suggests that RelB can act as both an activator and a repressor to regulate NF-κB-responsive gene expression. RelB also plays a dual regulatory role in targeting gene expression by recruiting co-activators or co-repressors [25]. In the present study, we first reported that RelB, an important transcription factor, represses the transcription of miR-193a-5p to mediate the phenotypic transition of VSMCs stimulated by Ang II.

In conclusion, these results suggest that inhibition of the RelB-miR-193a-5p axis to downregulate the expression of CCND1, CCNE1, and CXCR4 may offer new avenues for AA treatment.

## 5. Conclusion

In this study, we confirmed for the first time that the expression of the RelB-miR-193a-5p axis inhibits the proliferation and migration of VSMCs through CCND1, CCNE1, and CXCR4. This may provide new intervention targets for the prevention and treatment of AA.

## Author contribution

Yisi Liu and Chenghui Yan analyzed and interpreted the data. Xiaoxiang Tian, Dan Liu, Xiaolin Zhang, Chenghui Yan, and Yaling Han drafted and revised the manuscript.

## Data availability

Included in the article: The data that support the findings of this study are available in the methods and/or supplementary material of this article.

No new data: Data sharing does not apply to this article, as no datasets were generated or analyzed during the current study.

## Figure legends

**Figure S1.**
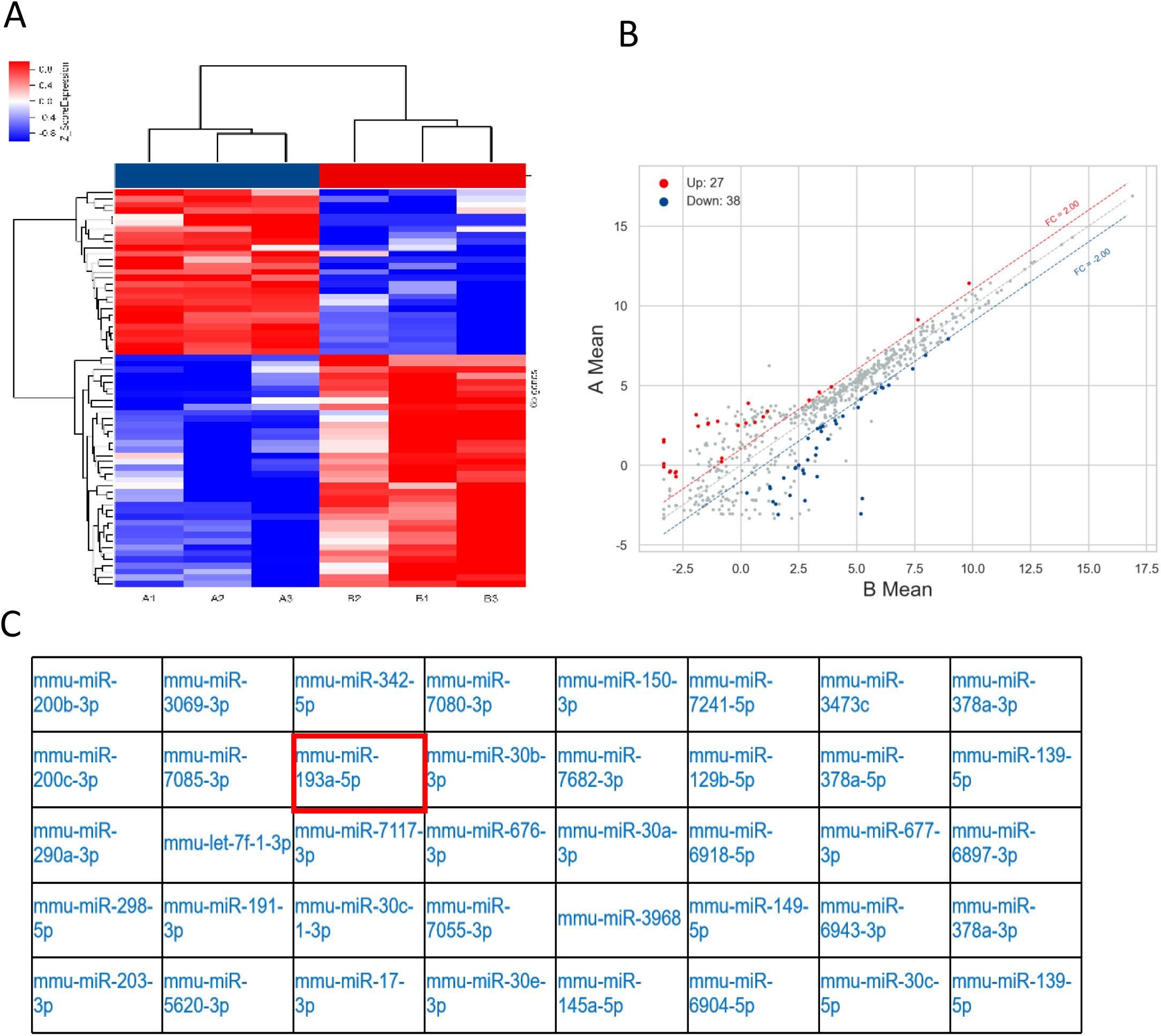
65 differentially expressed microRNAs in AA vascular tussue compared to control tissue in mice. A Cluster analysis chart(A), scatter plot(B) and list of down-regulated microRNAs(C).

**Figure S2.**
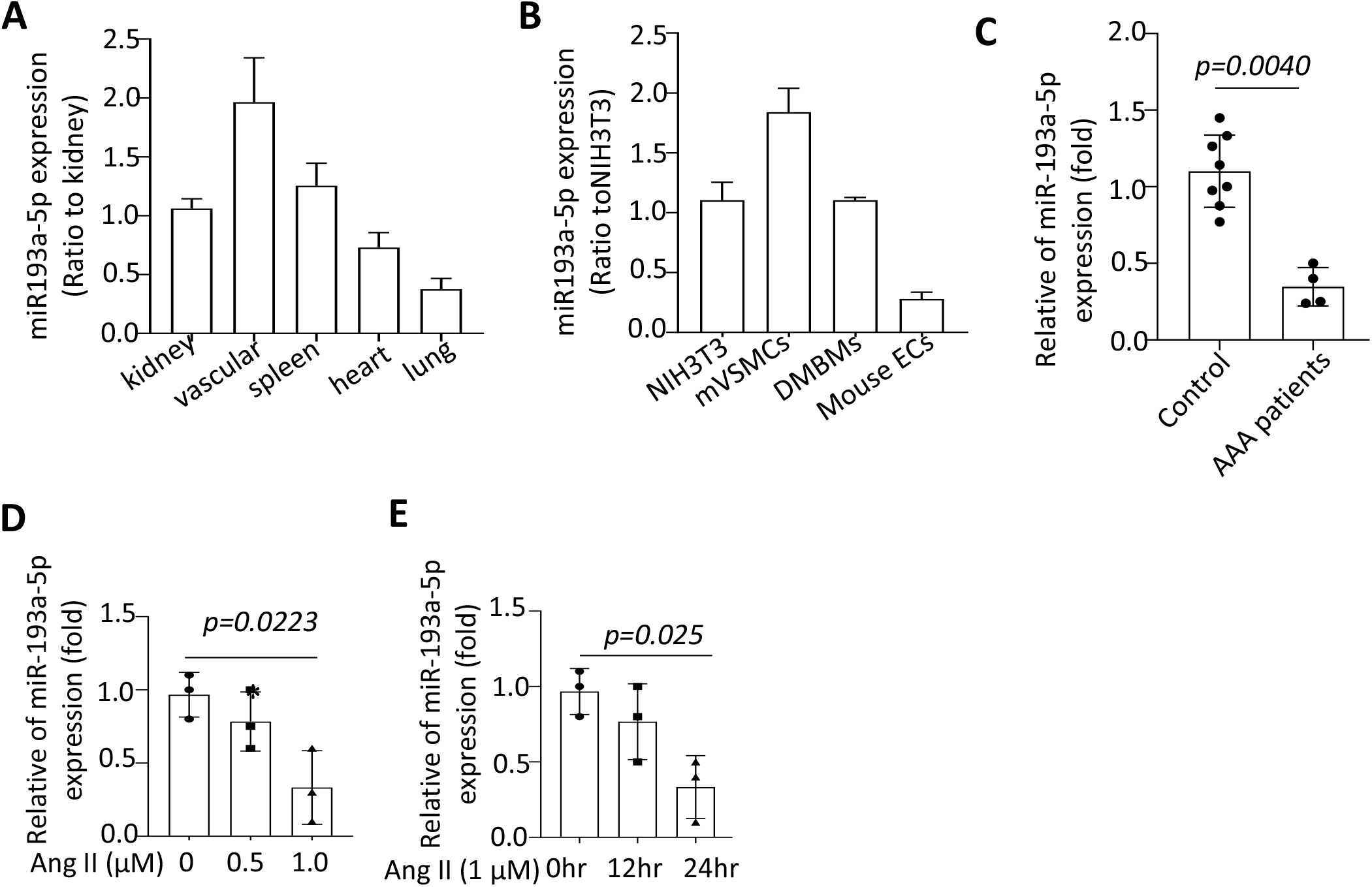
Expression levels of mir-193a-5p in different cell lines and pathological stimulation. (A) Real time quantitative PCR was detected the expression levels of mir-193a-5p in different mice tissue. (B) Real time quantitative PCR was detected the expression levels of mir-193a-5p in different cell lines from mice aortic tissue. (C) miR-193a-5p expression in serum from control group and AAA patient group. (D) miR-193a-5p expression in Ang II treated-VSMCs in dose-dependent manner. (E) miR-193a-5p expression in AngII treated-VSMCs in timedependent manner.

**Figure S3.**
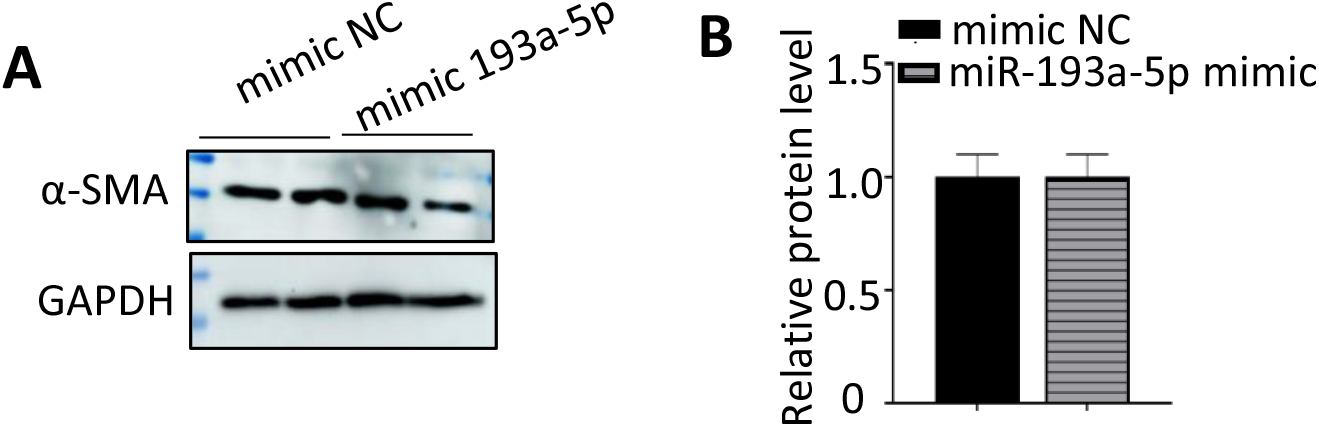
Expression of a-SMA in VSMCs with or without mimic mir-193a-5p transfection. Western blotting(A) and quantification analysis(B) detected the expression levels of a-SMA in VSMCs with or without miR-193a-5p transfection.

**Figure S4.**
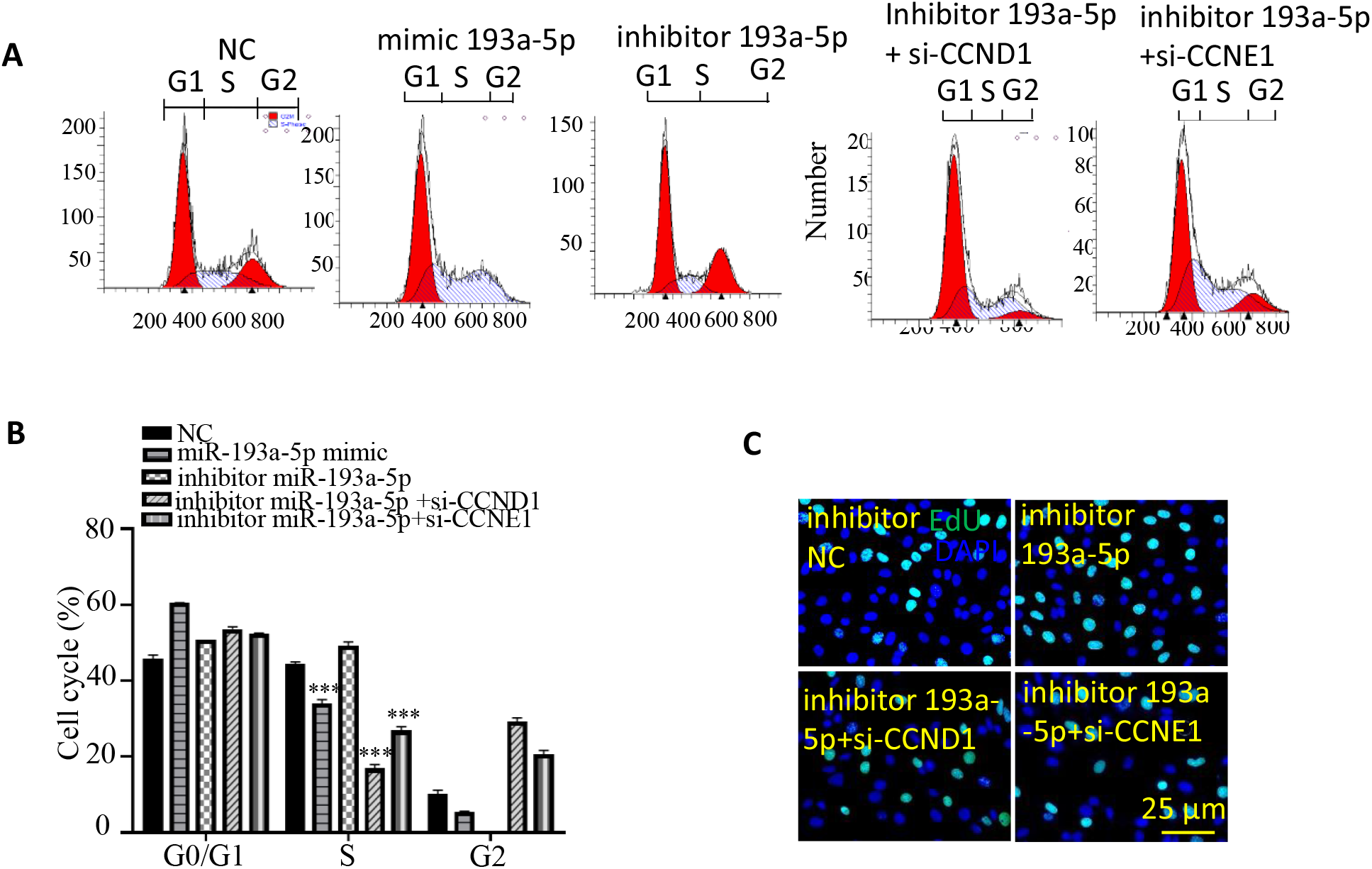
silence of CCND1 or CCNE1 inhibited the proliferative potential in VSMCs transfected with the miR-193a-5p inhibitor. (A) Flow cytometry was used to detect the effects of miR-193a-5p, CCND1 and CCNE1 on cell cycle. (B) Quanfication of the ratio of cell numbers distributed in different cell cycle. (C) EdU staining showed the proliferation of VSMCs when co-transfected inhibitor miR193a-5p and siCCND1 or siCCNE1 together.

**Figure S5.**
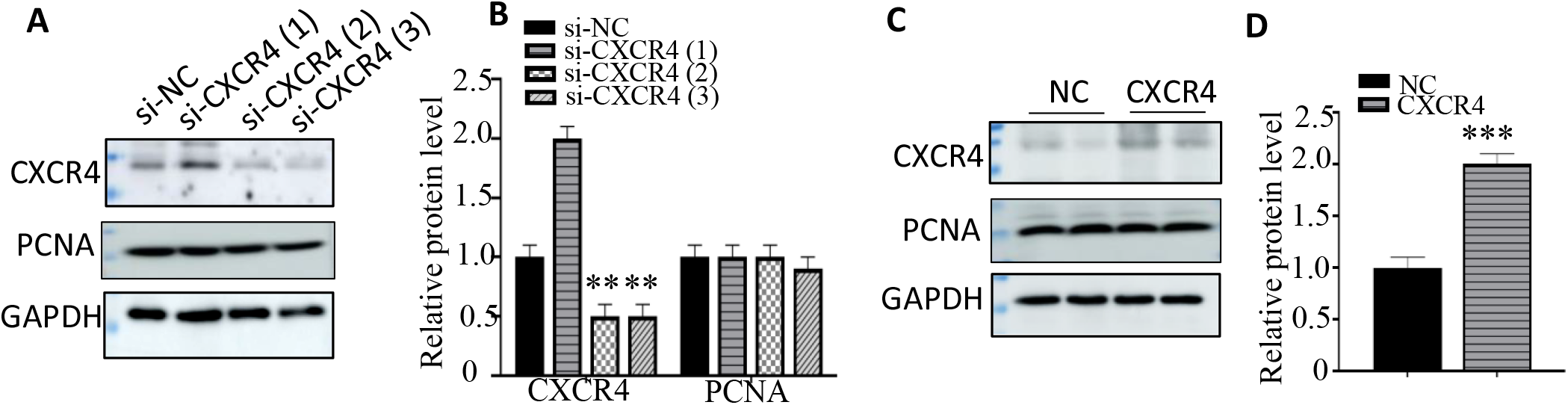
The expression of CXCR4 in VSMCs with siCXCR4 or pcDNA3-CXCR4 transfection. (A-B) The expression of CXCR4 was detected in VSMCs when transfected by siCXCR4 by Western blot. (C-D) Western blot was used to detect the expression of CXCR4 when VSMCs transfected the pcDNA3-CXCR4 plasmid.

**Figure S6.**
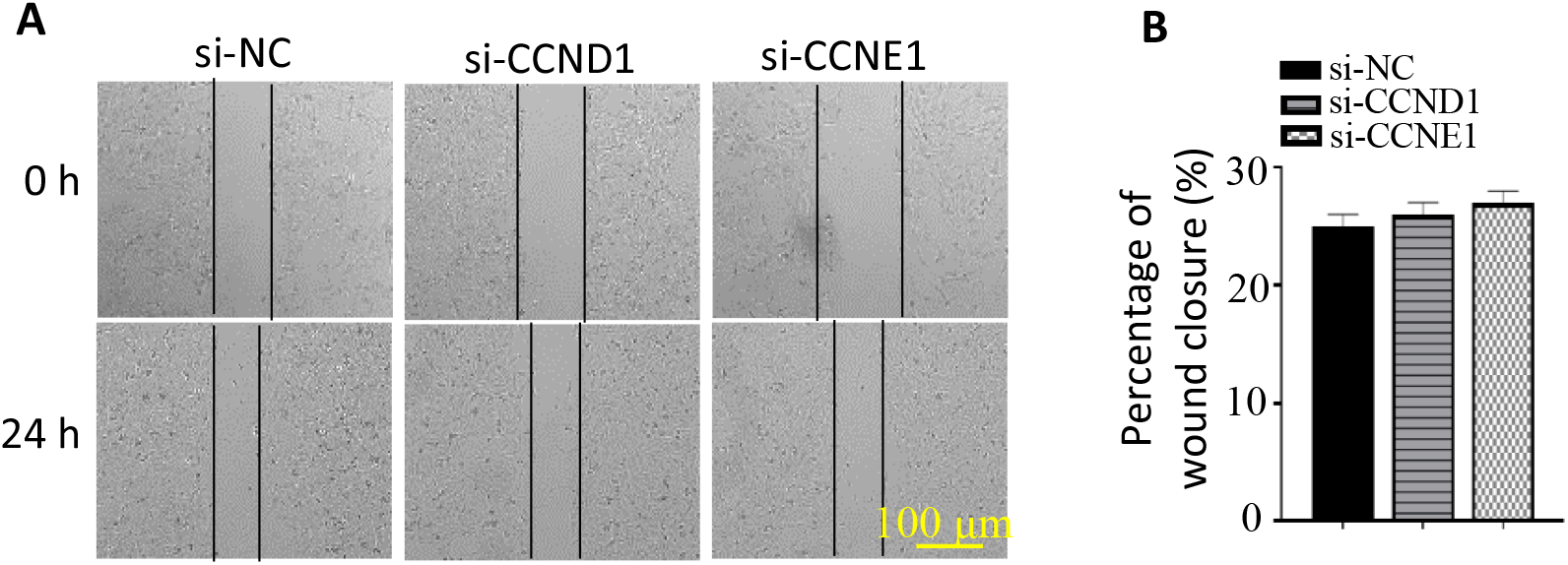
The effect of CCND1 and CCNE1 on migration. (A and B). The migration was observed by Wound healing assay and quantification.

**Major Resources Table.**

| **Compounds** | **Company** | **Catalog number** | **Address** |
| --- | --- | --- | --- |
| MOVAS | ATCC | Tings-98620 | Wanwu biocompany, hefei |
| Raw 264.7 and HEK293 | ATCC | TIB-71 | USA |
| Mice VECs | ATCC |  |  |
| FBS | Gibco | 10099141 | USA |
| DMEM | Gibco | 31331093 | USA |
| Angiotensin II (20 or 40 ng/ml) | Sigma | A9525 | USA |
| Mimic control, mimic miR193a-5p, inhibitor control, inhibitor miR-193a-5p | GENEWIZ INC |  | Suzhou, China |
| pcDNA3.1-CXCR4 , pcDNA3.1-CCNE1 , pcDNA3.1-CCND1 , pc.31-RelB, pc3.1-RelA and pcDNA3.1 empty vector | GENEWIZ, INC |  | Suzhou, China |
| si-RelB, si-CXCR4, si-CCNE1, si-CCND1 and si-HOXA5 | RIBOBIO, |  | Guangzhou, China |
| Dual-Luciferase Reporter assay system | Promega | E1910 | Beijing,China |
| RIPA | Thermo scientific | 89900 | Beijing,China |
| polyvinylidene fluoride membranes | Merck, Millipore | IPVH00010 | USA |
| TB Green Premix Ex Taq | Takara | Tli RNaseH Plus | Japan |
| CCK8 kit | Aibixin Company | abs50003 | Wuhan, China |
| Anti-PCNA, CCND1, CCNE1, a-SMA and CXCR4 | CST |  |  |
| Trizol reagent | Sigma-Aldrich | T9424 | USA |
| EdU kit | Ribo Bio |  |  |
| PI dye | BD Biosciences | 558662 | USA |

## References

1. Sakalihasan Natzi, Michel Jean-Baptiste, Katsargyris Athanasios et al. Abdominal aortic aneurysms [J]. Nat Rev Dis Primers, 2018, 4: 34.

2. Zhidong Zhang, Gangqiang Zou, Xiaosan Chen, et al. Knockdown of lncRNA PVT1 Inhibits Vascular Smooth Muscle Cell Apoptosis and Extracellular Matrix Disruption in a Murine Abdominal Aortic Aneurysm Model [J]. Mol Cells. 2019, 42(3):218–227

3. Rijan Gurung, Andrew-Mark Choong, Chin-Cheng Woo, et al. Genetic and Epigenetic Mechanisms Underlying Vascular Smooth Muscle Cell Phenotypic Modulation in Abdominal Aortic Aneurysm [J]. Int J Mol Sci. 2020, 21(17):6334.

4. Liu Ji-Ting, Liu Ze, Chen Yi et al. MicroRNA-29a Involvement in Phenotypic Transformation of Venous Smooth Muscle Cells Via Ten-Eleven Translocation Methylcytosinedioxygenase 1 in Response to Mechanical Cyclic Stretch [J]. J Biomech Eng, 2020, 142(5):051009

5. Chen L, Zheng S-Y, Yang C-Q et al. MiR-155-5p inhibits the proliferation and migration of VSMCs and HUVECs in atherosclerosis by targeting AKT1 [J]. Eur Rev Med Pharmacol Sci, 2019, 23:2223–2233.

6. Ji J, Xu Q, He X, Chen XL, Yang J. MicroRNA microarray analysis to detect biomarkers of aortic dissection from paraffin-embedded tissue samples [J]. Interact Cardiovasc Thorac Surg. 2020;31(2):239–247.

7. Moushi A, Michailidou K, Soteriou M, Cariolou M, Bashiardes E. MicroRNAs as possible biomarkers for screening of aortic aneurysms: a systematic review and validation study [J]. Biomarkers, 2018, 23(3):253–264.

8. Zhang W, Shang T, Huang C, Yu T, Liu C, Qiao T, Huang D, Liu Z, Liu C. Plasma microRNAs serve as potential biomarkers for abdominal aortic aneurysm [J]. Clin Biochem, 2015, 48(15):988–992.

9. Noorolyai S, Baghbani E, Shanehbandi D, et al. miR-146a-5p and miR-193a-5p Synergistically Inhibited the Proliferation of Human Colorectal Cancer Cells (HT-29 cell line) through ERK Signaling Pathway [J]. Adv Pharm Bull. 2021 Sep; 11(4):755–764.

10. Zhang Y, Jiang F, He H, et al. Identification of a novel microRNA-mRNA regulatory biomodule in human prostate cancer[J]. Cell Death Dis. 2018 Feb 21;9(3):301.

11. Jun Ma, Li-Na Han, Jian-Rui Song, et al. Long noncoding RNA LINC01234 silencing exerts an anti-oncogenic effect in esophageal cancer cells through microRNA-193a-5p-mediated CCNE1 downregulation[J]. Cell Oncol (Dordr). 2020;43(3):377–394.

12. Ullery BW, Hallett RL, Fleischmann D. Epidemiology and contemporary management of abdominal aortic aneurysms [J]. Abdom Radiol (NY), 2018, 43(5):1032–1043.

13. Wang YD, Liu ZJ, Ren J, Xiang MX. Pharmacological Therapy of Abdominal Aortic Aneurysm: An Update [J]. Curr Vasc Pharmacol, 2018, 16(2):114–124.

14. Davis FM, Rateri DL, Daugherty A. Abdominal aortic aneurysm: novel mechanisms and therapies [J]. Curr Opin Cardiol, 2015, 30(6):566–573.

15. Pedroza AJ, Tashima Y, Shad R, et al. Single-Cell Transcriptomic Profiling of Vascular Smooth Muscle Cell Phenotype Modulation in Marfan Syndrome Aortic Aneurysm. Arterioscler Thromb Vasc Biol. 2020;40(9):2195–2211.

16. Rombouts KB, van Merrienboer TAR, Ket JCF, et al. The role of vascular smooth muscle cells in the development of aortic aneurysms and dissections. Eur J Clin Invest. 2022;52(4):e13697.

17. Joviliano Edwaldo Edner, Ribeiro Mauricio Serra, Tenorio Emanuel Junior Ramos, MicroRNAs and Current Concepts on the Pathogenesis of Abdominal Aortic Aneurysm [J]. Braz J Cardiovasc Surg, 2017, 32:215–224.

18. Chou Nan-Hua, Lo Yi-Hao, Wang Kuo-Chiang et al. MiR-193a-5p and −3p Play a Distinct Role in Gastric Cancer: miR-193a-3p Suppresses Gastric Cancer Cell Growth by Targeting ETS1 and CCND1 [J]. Anticancer Res, 2018, 38:3309–3318.

19. Azar Mohammad Reza Mohammad Hoseini, Aghazadeh Hamed, Mohammed Halgurd Nadhim et al. miR-193a-5p as a promising therapeutic candidate in colorectal cancer by reducing 5-FU and Oxaliplatin chemoresistance by targeting CXCR4 [J]. Int Immunopharmacol, 2021, 92: 107355.

20. Xie Mei, Zhao Fen, Zou Xiaoling et al. The association between CCND1 G870A polymorphism and colorectal cancer risk: A meta-analysis [J]. Medicine (Baltimore), 2017, 96:e8269.

21. Gorski Justin W, Ueland Frederick R, Kolesar Jill M, CCNE1 Amplification as a Predictive Biomarker of Chemotherapy Resistance in Epithelial Ovarian Cancer [J]. Diagnostics (Basel), 2020, 10(5):279.

22. Zielińska Karolina A, Katanaev Vladimir L, The Signaling Duo CXCL12 and CXCR4: Chemokine Fuel for Breast Cancer Tumorigenesis [J]. Cancers (Basel), 2020, 12(10):3071.

23. Laura Martínez-Muñoz, José Miguel Rodríguez-Frade, Rubén Barroso, et al. Separating Actin-Dependent Chemokine Receptor Nanoclustering from Dimerization Indicates a Role for Clustering in CXCR4 Signaling and Function [J]. Mol Cell. 2018;70(1):106–119.

24. Nenasheva VV, Kovaleva GV, Uryvaev LV, et al. Enhanced expression of trim14 gene suppressed Sindbis virus reproduction and modulated the transcription of a large number of genes of innate immunity. Immunol Res[J]. 2015 Jul;62(3):255–62.

25. Yang MG, Sun L, Han J, Zheng C, Liang H, Zhu J, Jin T. Biological characteristics of transcription factor RelB in different immune cell types: implications for the treatment of multiple sclerosis. Mol Brain[J]. 2019, 12:115.

